# Fundamental Differences: A Basis Set for Characterizing Inter-Individual Variation in Resting State Connectomes

**DOI:** 10.1101/326082

**Authors:** Chandra Sripada, Mike Angstadt, Saige Rutherford, Daniel Kessler, Yura Kim, Mike Yee, Liza Levina

## Abstract

Resting state functional connectomes are massive and complex. It is an open question, however, whether connectomes differ across individuals in a correspondingly massive number of ways, or whether most differences take a small number of characteristic forms. We systematically investigated this question and found clear evidence of low-rank structure in which a modest number of connectomic components, around 50-150, account for a sizable portion of inter-individual connectomic variation. This number was convergently arrived at with multiple methods including estimation of intrinsic dimensionality and assessment of reconstruction of out-of-sample data. We demonstrate that these connectomic components enable prediction of a broad array of neurocognitive and clinical variables. In addition, using stochastic block modeling-based methods, we show these components exhibit extensive community structure reflecting interrelationships between intrinsic connectivity networks. We propose that these connectivity components form an effective basis set for quantifying and interpreting inter-individual connectomic differences, and for predicting behavioral/clinical phenotypes.

## 1. Introduction

Resting state functional connectomics has emerged as a leading method for mapping the organization of human brain networks (Biswal *et al.*, 2010; Van Dijk *et al.*, 2010; Buckner, Krienen and Yeo, 2013; Lee, Smyser and Shimony, 2013; Smith *et al.*, 2013). In addition, it presents a major opportunity for elucidation of the brain basis of individual differences (Barch, 2013; Castellanos *et al.*, 2013; Matthews and Hampshire, 2016; Menon, 2011): functional networks are thought to be critical substrates for major neurocognitive and behavioral phenotypes (Laird *et al.*, 2011; Bressler and Menon, 2010; Mattar *et al.*, 2015), so across-individual differences in network organizations may predict differences in these phenotypes (Kelly *et al.*, 2012; Dubois and Adolphs, 2016). The eventual goal is to refine phenotypic prediction sufficiently that functional connectomes can serve as reliable, objective “biomarkers” of clinically meaningful traits and dimensions (Castellanos *et al.*, 2013; Kaiser, 2013; Woo *et al.*, 2017; Bassett, Xia and Satterthwaite, 2018; Satterthwaite, Xia and Bassett, 2018).

Notably, while attempts to utilize functional connectomes for prediction of individual differences are numerous (Kelly *et al.*, 2012; Dubois and Adolphs, 2016), attempts to descriptively assess the nature, kind, and extent of population-wide inter-individual functional connectomic variation remain scarce, c.f., Mueller *et al.*, 2013; Gordon *et al.*, 2017. One important open question concerns the dimensionality of inter-individual variation.

In high dimensional data, there is often substantial dependency in the feature set, and it is often useful for a wide of variety of purposes—computation, interpretation, explanation, and prediction—to identify low-rank structure in the data, i.e major components that explain a substantial portion of the variation. Over the last 15 years, there has been extensive work in detecting low-rank structure in *intra-individual* across-time variation in the connectome, i.e the tendency of distributed brain regions to exhibit coherent fluctuations in their BOLD time series (Greicius *et al.*, 2004; van de Ven *et al.*, 2004; Beckmann *et al.*, 2005). This work has culminated in the identification of a small number of intrinsic connectivity networks (ICNs) as major components of intra-individual cross-time variation (Power *et al.*, 2011; Yeo *et al.*, 2011; Buckner, Krienen and Yeo, 2013). These networks, in turn, have played central roles in recent models and explanations of cognitive capacities and behavioral phenotypes (Bressler and Menon, 2010; Laird *et al.*, 2011; Menon, 2011; Barch, 2013; Cole *et al.*, 2013; Mattar *et al.*, 2015).

Importantly, however, there have not been corresponding systematic attempts to identify low-rank structure in patterns of *inter-individual* variation (but see Kessler *et al.*, 2014; Kessler, Angstadt and Sripada, 2016; Amico *et al.*, 2017 for limited attempts). This is the question we address in this study. That is, analogous to the intra-individual case, are there major components of inter-individual variation that explain a sizable portion of cross-individual connectomic differences, and that can be effectively harnessed for the purposes of understanding and predicting phenotypes of interest?

In this study, we provide evidence that the answer to this question is yes. Using convergent methods, we show that a modest number of connectivity components, around 50-150, do indeed capture a sizable share of inter-individual differences, and they together constitute a highly effective basis set for phenotypic prediction. Thus, while the resting state connectome is a massive and complex object encompassing tens to hundreds of thousands of connections (depending on the parcellation), differences in a fairly small set of components explain a sizable portion of how any two individuals meaningfully differ.

## 2. Methods

### 2.1 Subjects and Data Acquisition

All subjects and data were from the HCP-1200 release (Van Essen *et al.*, 2013; WU-Minn HCP, 2017). All subjects provided informed consent. Subject recruitment procedures and informed consent forms, including consent to share de-identified data, were approved by the Washington University institutional review board. Four runs of resting state fMRI data (14.5 minutes each; two runs per day over two days) were acquired on a modified Siemans Skyra 3T scanner using multiband gradient-echo EPI (TR=720ms, TE=33ms, flip angle = 52°, multiband acceleration factor = 8, 2mm isotropic voxels, FOV = 208×180mm, 72 slices, alternating RL/LR phase encode direction). T1 weighted scans were acquired with 3D MPRAGE sequence (TR=2400ms, TE=2.14ms, TI=1000ms, flip angle = 8, 0.7mm isotropic voxels, FOV=224mm, 256 sagittal slices). T2 weighted scans were acquired with a Siemens SPACE sequence (TR=3200ms, TE=565ms, 0.7mm isotropic voxels, FOV=224mm, 256 sagittal slices).

Subjects were eligible to be included if they had structural T1 and T2 data and had 4 complete resting state fMRI runs (14m 30s each; 1206 subjects total in release files, 1003 with full resting state and structural).

### 2.2 Data Preprocessing

Processed volumetric data from the HCP minimal preprocessing pipeline including ICA-FIX denoising were used. Full details of these steps can be found in Glasser (2013) and Salimi-Korshidi (2014). Briefly, T1w and T2w data were corrected for gradient-nonlinearity and readout distortions, inhomogeneity corrected, and registered linearly and non-linearly to MNI space using FSL’s FLIRT and FNIRT. BOLD rfMRI data were also gradient-nonlinearity distortion corrected, rigidly realigned to adjust for motion, fieldmap corrected, aligned to the structural images, and then registered to MNI space with the nonlinear warping calculated from the structural images. Then FIX was applied on the data to identify and remove motion and other artifacts in the timeseries. These files were used as a baseline for further processing and analysis (e.g., MNINonLinear/Results/rfMRI_REST1_RL/rfMRI_REST1_RL_hp2000_clean.nii.gz from released HCP data).

Images were smoothed with a 6mm FWHM Gaussian kernel, and then resampled to 3mm isotropic resolution. This step as well as the use of the volumetric data, rather than the surface data, were done to allow comparability with other large datasets in ongoing and planned analyses that are not amenable to surface-based processing.

The smoothed images then went through a number of resting state processing steps, including a motion artifact removal steps comparable to the type B (i.e., recommended) stream of Siegel et al. (2017). These steps include linear detrending, CompCor to extract and regress out the top 5 principal components of white matter and CSF (Behzadi *et al.*, 2007), bandpass filtering from 0.1-0.01Hz, and motion scrubbing of frames that exceed a framewise displacement of 0.5mm. Subjects with more than 10% of frames censored were excluded from further analysis, leaving 966 subjects. A resting state quality control plot (Power *et al.*, 2014) relating motion effects by edge length showed a near zero mean (0.006), low dispersion around the mean (sd 0.06) and absence of a meaningful distance-dependent relationship.

### 2.3 Connectome Generation

We next calculated spatially-averaged time series for each of 264 4.24mm radius ROIs from the parcellation of Power et al. (Power *et al.*, 2011). We then calculated Pearson’s correlation coefficients between each ROI. These were then were transformed using Fisher’s r to z-transformation.

### 2.4 Train/Test/Retest Split

The 966 subjects after exclusions were divided into three groups. First, 38 subjects who had two separate completed scans were pulled aside for later test-retest reliability analysis. Of the remaining subjects, 18 did not have complete behavioral data for our analyses so were excluded. Next, 100 unrelated subjects were randomly selected from all unrelated subjects to serve as our held out test set, with the other 810 serving as our training set.

### 2.5 Estimation of Intrinsic Dimensionality

In the training dataset, each subject’s connectome was vectorized and concatenated yielding an 810 subjects × 34,716 connections matrix. We estimated the number of intrinsic dimensions of this matrix using two methods.

First, we used a maximum likelihood estimation method based on distance between close neighbors (Levina and Bickel, 2004), appropriate for low-dimensional data that is embedded in a high-dimensional space in a complicated, potentially non-linear, fashion. Levina and Bickel (2004) provides a full derivation of the estimator using a Poisson approximation and demonstrates improved performance relative to alternatives in simulated and real data. The method averages over a range of values of *k*, the number of nearest neighbors, from *k_1_* to *k_2_*. We used the default values *k_1_* = 10 to *k_2_* = 20 suggested by the original analysis.

Second, we used the method of Choi et al. (2017), which attempts to calculate an upper bound on the on the number of dimensions with exact type 1 error control. This is a distribution-based method that leverages a post-selection inference framework, and extends the work of Taylor, Loftus, and Tibshirani (2016) to the PCA setting.

To visualize the presence of low-rank structure, we ordered the components by eigenvalue (i.e., percent variance explained), and plotted these eigenvalues. Next we constructed a null distribution of eigenvalues by permutation methods. Specifically, we permuted columns of the data matrix separately for each subject. We plotted the permutation mean and 95% confidence interval for the null distribution.

### 2.6 Principal Component Analysis

The subjects *x* connections matrix from the training dataset was next submitted to principal components analysis using the pca function in MATLAB, yielding 809 components ordered by descending eigenvalues..

### 2.7 Assessing Out-of-Sample Reconstruction

We examined the ability of an *n*-sized basis set (consisting of the first *n* PCA components ordered by descending eigenvalues), to reconstruct out-of-sample data, systematically varying the size of *n*. First, a full set of 809 PCA components were learned on the training dataset. Next, for each value of *n* from 1 to 809, we did the following: Using multiple regression, each subject in the test dataset was reconstructed as linear combination of the components of an *n*-sized basis set. Goodness of reconstruction was measured by calculating the Pearson’s correlation across edges between actual versus reconstructed connectomes for each subject, and averaging across subjects.

### 2.8 Assessing Phenotypic Prediction

#### 2.8.1 HCP Phenotypic Measures

We used a total of 11 phenotypes from the HCP data. Factor analysis, implemented in SPSS 23 (IBM, Armonk, NY), was used to produce two neuropsychological factors from the HCP task data. First, a general executive factor was created based on overall accuracy for three tasks: *n*-back working memory task, relational processing task, and Penn Progressive Matrices task. Factor loadings were 0.81, 0.80, and 0.76 respectively, and the factor accounted for 62.2% of the variance in the variables. A speed of processing variable was created based on three NIH toolbox tasks: processing speed, flanker task, and card sort task (all age-adjusted performance), similar to Carlozzi *et al.*, 2015. Of note, the first of these three tasks is designed to be a measure of processing speed, while the latter two primarily reflect processing speed because for most subjects in the HCP dataset, accuracy is close to ceiling (Slotkin *et al.*, 2012). This variable had loadings of 0.75, 0.81, and 0.82 respectively, and the factor accounted for 63.0% of the variance in the variables. From the Adult Self Report (ASR) instrument (Achenbach, 2009), we used three scale-derived summary scores for psychopathology: overall internalizing, overall externalizing, and attention. In addition, from the Neuroticism/Extroversion/Openness Five Factor Inventory instrument (McCrae and Costa, 2004), we used the five personality factors: openness to experience, conscientiousness, extroversion, agreeableness, and neuroticism. Finally, we used the Penn Progressive Matrices task by itself as it has been featured in other connectome-based prediction studies of HCP data (Finn *et al.*, 2015; Ma, Guntupalli and Haxby, 2017).

In an additional analysis, we used multiple regression to remove a number of potential confounds from each of the 11 phenotypic variables. Variables regressed from the phenotypes were: age, age2, mean FD, mean FD2, gender, brain size (S BrainSeg Vol), brain size2, and multiband reconstruction algorithm version number (fMRI 3T ReconVrs). Analyses involving phenotypic prediction (§2.8.3 and §2.8.4) were then repeated with the confound-cleansed phenotypes. Results were broadly similar to the original analyses, and are presented in the Supplement.

#### 2.8.2 Brain Basis Set Modeling

To generate predictions of phenotypes from a basis set consisting of *n* components, we used Brain Basis Set (BBS) modeling, similar to the approach introduced in Kessler, Angstadt, and Sripada 2016. In a training dataset, we calculate the expression scores for each of the *n* components for each subject. We then fit a linear regression model with these expression scores as predictors and the phenotype of interest as the outcome, saving **B**, the *n × 1* vector of fitted coefficients, for later use. In a test dataset, we again calculate the expression scores for each of the *n* components for each subject. Our predicted phenotype for each test subject is the dot product of **B** learned from the training dataset with the vector of component expression scores for that subject.

#### 2.8.3 10-fold cross validation procedure

We assessed prediction of HCP phenotypes as a function of number of components in the predictive basis set, in order to identify the presence of plateaus where adding additional components does not enhance predictive accuracy. This analysis was performed using a 10-fold cross-validation procedure within the training dataset split described above (to preserve the test dataset for additional analyses described below). On each of the ten folds, we used the training partition to learn new PCA components and then fit beta coefficients for BBS modeling. We then made predictions for the phenotypes in the held out test partition. The correlations between actual phenotype and predicted phenotype were then averaged across the ten folds.

#### 2.8.4 Comparison with CPM

To further assess the effectiveness of a low-rank basis set for capturing phenotypic differences in the HCP dataset, we compared the accuracy of phenotypic predictions derived from the 100 component basis set (coupled with CBS modeling) with predictions derived from an alternative leading method: connectome predictive modeling (CPM) (Shen *et al.*, 2017), which has achieved excellent results in a number of studies using diverse phenotypes (Finn *et al.*, 2015; Rosenberg *et al.*, 2016; Yoo *et al.*, 2018; Beaty *et al.*, 2018; Lake *et al.*, 2018). In brief, CPM is first trained with every edge of the connectome to identify edges that are predictive of the phenotype of interest above some prespecified level (e.g., Pearson’s correlation with significance of *p*< 0.01). The sum of weights for these specified edges is then calculated for each test subject, and these sums serve as “predicted scores” that are correlated with the actual phenotypic scores. CPM treats positively and negatively predictive edges differently, and we focus on the positive edges in the main article, following the typical practice of its authors, and present results for negative edges in the Supplement.

### 2.9 Density of Parcellation Analysis

To assess the robustness of the analysis to parcellations of systematically varying densities, we used the set of parcellations created by Craddock et al. (2011). These parcellations (available here: http://ccraddock.github.io/cluster_roi/atlases.html) were produced with a spatially constrained spectral clustering approach that, for preset values of *K*, produces approximately *K* functionally and spatially coherent regions. We utilized parcellations with *K* ranging from 100-900 in intervals of 100. For each parcellation, we repeated steps 3 through 8 of the above analysis in order to assess whether our three methods for identifying low-rank structure (assessment of: intrinsic dimensionality, out-of-sample-reconstruction, and phenotypic prediction) differed according to parcellation density. Of note, our implementation of the method of Choi et al. did not converge for larger parcellations (K > 500) and so we focus on the the method of Levina and Bickel for this analysis.

### 2.10 Assessing Community Structure

For all 809 components, we assessed the presence of community structure corresponding to ICNs from the parcellation of Power et al. (2011) using a stochastic block model (SBM; Holland, Laskey and Leinhardt, 1983), a well established generative model for graphs, coupled with a non-parametric testing procedure. For each of the 809 components, we first fix node community assignments according to the Power parcellation (Power *et al.*, 2011), and then estimate the parameters of a SBM with these fixed assignments. We replace the Bernoulli distribution assumption on binary edges made by the classical SBM with a normal distribution assumption on edge weights, since we work with Fisher-transformed correlations as edge weights. Once these parameters are estimated, we summarize the fit with the profile log-likelihood statistic. We then randomly permute node labels many times, keeping the total number of nodes in each of the communities fixed, and obtain a profile likelihood value from each of these fits corresponding to permuted node labels. We then obtain a p-value by comparing the profile likelihood for the Power parcellation to the empirical null distribution of profile likelihoods. Finally, the p-values for the 809 components were adjusted for multiple comparisons using Bonferroni’s correction, to control the Family-Wise Error Rate at α = 0.05. A more detailed description of this procedure is provided in the Supplement.

### 2.11 Test/Retest Reliability

Test-retest reliability was assessed in 38 subjects in the HCP test-retest dataset. Reliability was assessed with intra-class correlation (ICC) statistic, specifically type (2,1) according to the scheme of Shrout and Fleiss (1979). For each subject, ICC’s were calculated for each individual edge as well as for expression scores for each component in the 100-member basis set. Since aggregating edges can itself improve ICC, we also examined ICC’s for “random” aggregations of edges created by permuting the edges of each of the 100 components. For each component, 1000 randomly permuted components were created in this way, and ICC’s for the expressions of these components were calculated.

## 3 Results

### 3.1 There is convergent evidence for substantial low-rank structure in cross-individual connectomic variation based on three different methods

#### 3.1.1 Method 1: Assessing intrinsic dimensionality

Figure 1 shows the percent variance explained (i.e., eigenvalues) for all 809 components (in blue). Also plotted is mean eigenvalues for 1000 realizations of random data created through permutation methods (in red). This plot provides initial suggestive evidence of significant low-rank stricture in the data, indicated by the substantially elevated variance explained by early components derived from observed connectomes relative to what components derived from random data.

**Figure 1:**
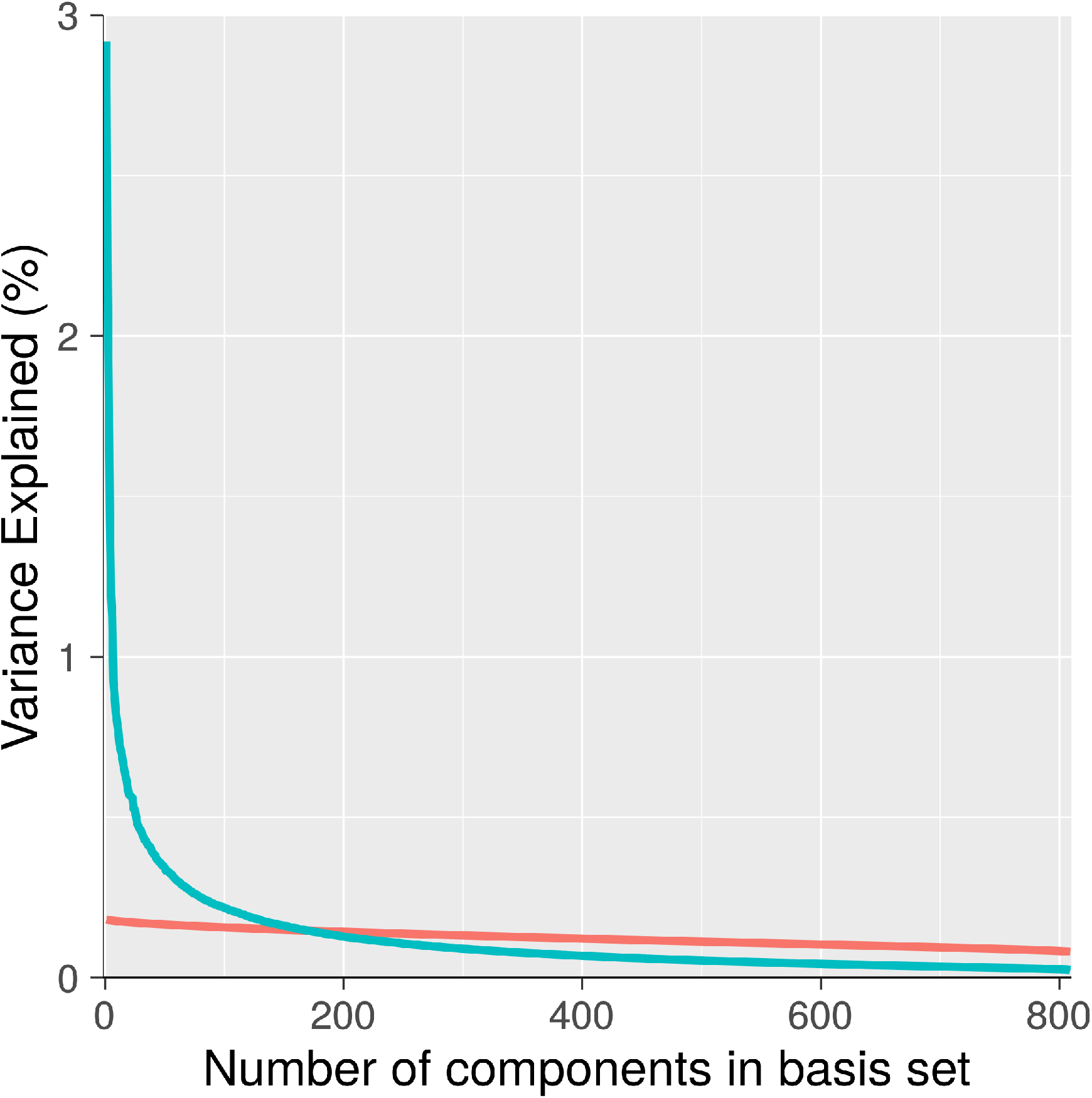
Percent Variance Explained For Components Derived From Actual Versus Random Data. For observed components (blue), percent variance explained of early components is much larger than mean percent variance explained of components derived from random data (red). This pattern is suggestive of substantial low-rank structure in observed connectomes.

We next turned to quantitative dimensionality estimation procedures. Applying the maximum likelihood method from Levina and Bickel (2004) yielded an estimated dimensionality of 62. Applying the dimensionality estimation method of Choi et al (2017) found an upper bound of 147 components with *alpha* set at 0.05. Importantly, these two results should be seen as complementary and not necessarily in tension, as the Levina and Bickel method attempts to arrive at the number of components that is *most likely* given the data, while the Choi et al method attempts to provide an *upper bound* on the number of components, with statistical control over type 1 errors. Taken together, these methods provide strong initial evidence for substantial low-rank structure in cross-individual connectomic variation. In addition, they suggest a plausible range for the number of true dimensions in the data as being somewhere between 50 and 150.

#### 3.1.2 Method 2: Assessing out of sample reconstruction

A second method for detecting and quantifying low-rank structure relies on examining the ability of the PCA components to accurately reconstruct connectomes from an independent test sample, i.e a sample that was not used to generate the components. Figure 2 shows the Pearson’s correlations between actual test sample connectomes and connectomes reconstructed with a PCA-derived basis set, as a function of the number of components in the basis set. Using all 809 components in the basis set, this correlation was 0.68, and this represents the ceiling correlation that is achievable. With 50, 100, and 150 components, the correlation is 0.47, 0.53, 0.57, respectively. This represents, respectively, 69%, 78%, and 84% of ceiling, and it provides additional evidence that a low-rank representation captures a sizable portion of the generalizable variance in the data.

**Figure 2:**
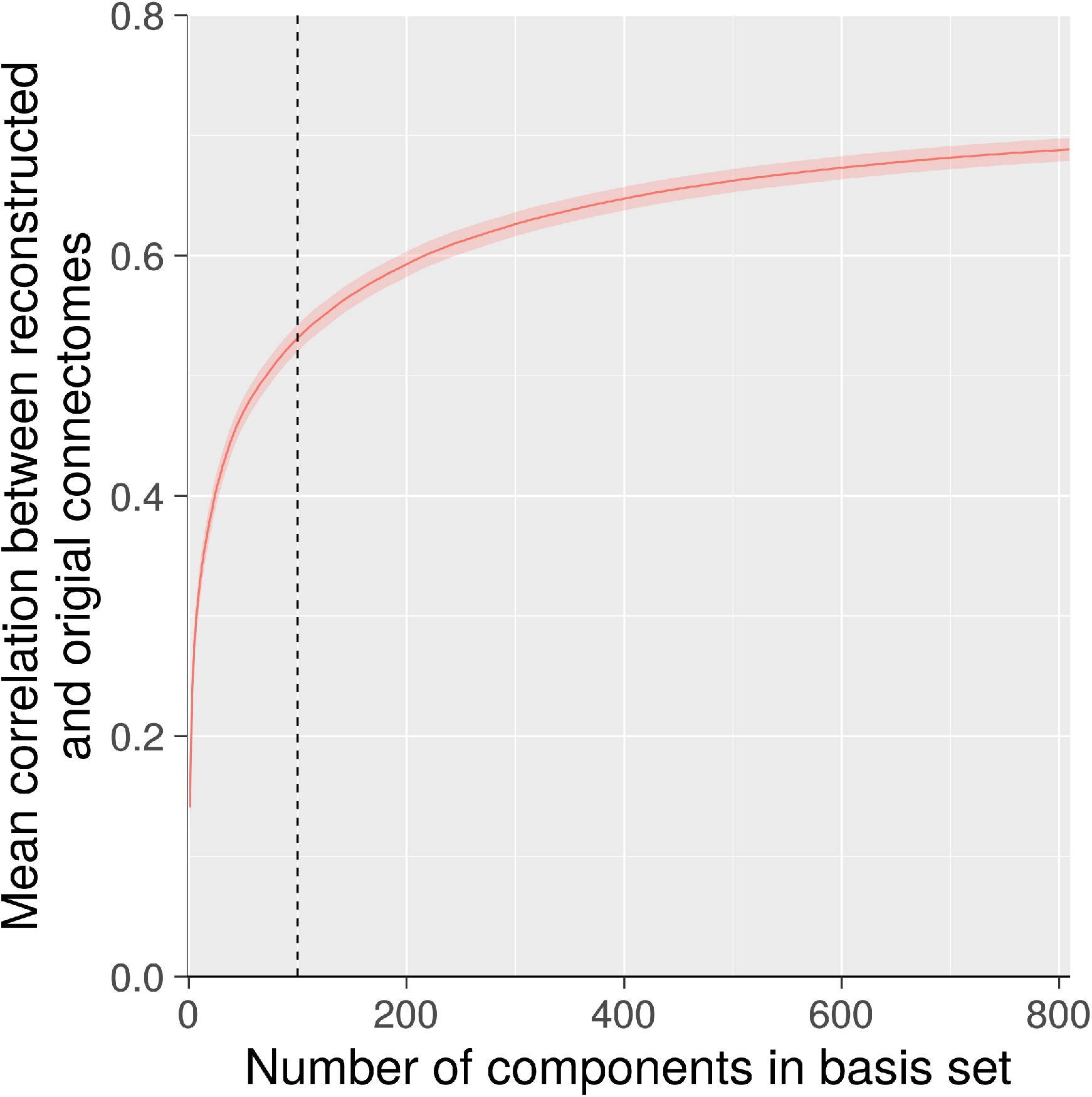
Out of Sample Reconstruction of Connectomes. With 100 components (dashed line), the correlation between actual and reconstructed connectomes is 0.50. Importantly, this correlation is only 0.68 using all 809 components, so a basis set consisting of 100 components achieves roughly three fourths of the “ceiling” correlation that is achievable.

#### 3.1.3 Method 3: Assessing predictive accuracy with respect to a broad range of HCP phenotypes

An additional means to assess low-rank structure consists in examining prediction of criterion variables: If a modest sized basis set captures a large portion of cross-individual variation, then it ought to predict a broad range of behavioral and clinical phenotypes (that are plausibly linked to functional connectomic variation) similarly to the full unreduced dataset.

For each of 11 phenotypes, we used the BBS modeling method to make predictions of phenotype values for each subject based on connectomic component expression scores. We applied BBS to the 810 subjects in the training dataset in a 10-fold cross validation procedure (see Methods, §8.2). As shown in Figure 3, there is a noticeable plateau at around 50-100 components for most of the phenotypes: Adding further components to the basis set beyond this number does not appreciably increase accuracy of phenotypic prediction. Table S1 shows the correlations between predicted and actual phenotypes across three basis set sizes: 50, 100, and 150 components. All three basis sets perform similarly, though there is a slight advantage for the 100-component basis set, especially with regard to the processing speed factor.

**Figure 3:**
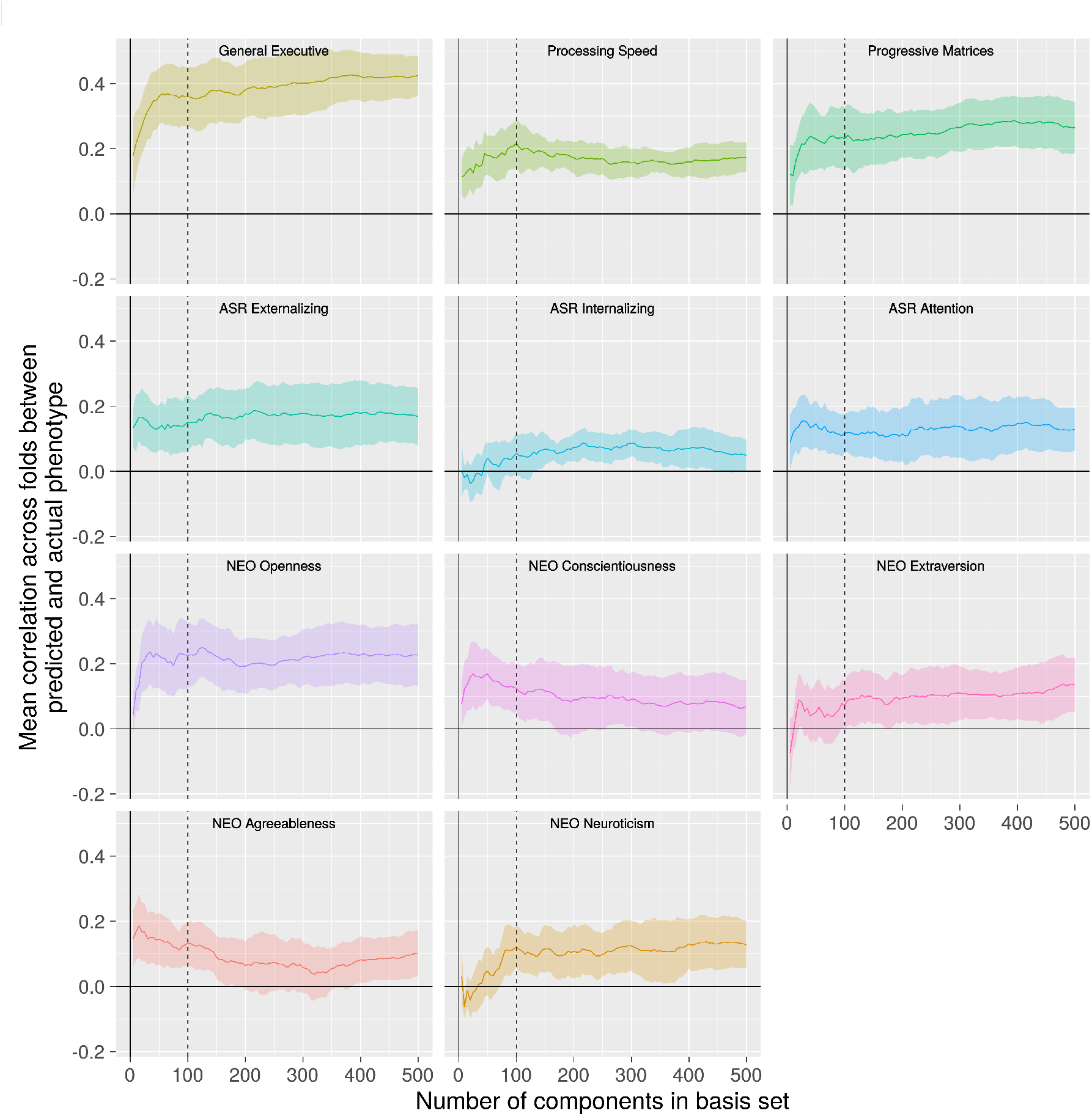
Phenotype Predictive Accuracy as a Function of Basis Set Size. For most phenotypes, there is a plateau after 50 to 100 components (dotted line) after which adding further components to the prediction model basis set does not appreciably improve performance.

**Figure 4:**
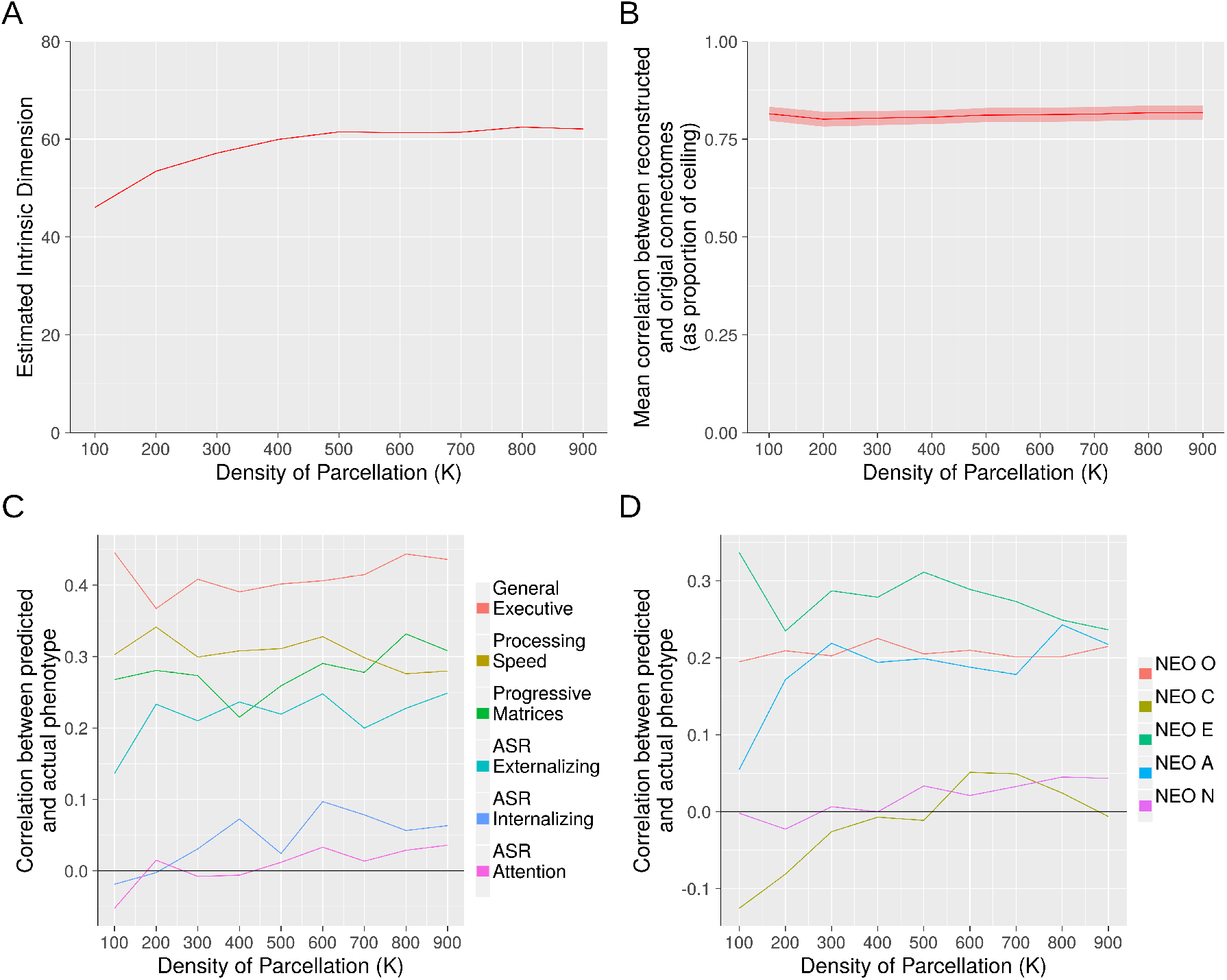
Assessing Role of Parcellation Density. Three methods for identifying low-rank structure yielded stable results across parcellations of varying density. *Panel A*: Estimation of intrinsic dimensionality with the method of Levina and Bickel, 2004. *Panel B*: Out-of-sample reconstruction. *Panels C and D*: Predictive accuracy with respect to 11 HCP phenotypes. Panels B through D used a 100-component basis set.

**Table 1:** Pearson’s correlations between actual and predicted phenotypes for two different predictive modeling approaches. *BBS = Brain Basis Set Modeling (with 100 component basis set); CPM = Connectome Predictive Modeling* (Shen *et al.*, 2017). *=statistically significant difference at *p*<0.05.

| ***Phenotype*** | **BBS** | **CPM** |
| --- | --- | --- |
| *General Executive* | 0.44 | 0.42 |
| *Processing Speed* | 0.39* | 0.23 |
| *Penn Progressive Matrices* | 0.30 | 0.32 |
| *ASR Externalizing* | 0.24* | 0.03 |
| *ASR Internalizing* | 0.20 | 0.04 |
| *ASR Attention* | 0.15* | 0.00 |
| *NEO-Openness* | 0.18 | 0.11 |
| *NEO-Conscientiousness* | 0.19 | 0.15 |
| *NEO-Extroversion* | 0.13 | 0.04 |
| *NEO-Agreeableness* | 0.19 | 0.10 |
| *NEO-Neuroticism* | 0.00 | 0.05 |

To further assess the performance of a modest sized basis set in predicting phenotypes of interest, we compared performance with CPM, a leading alternative method for phenotypic prediction that is trained on the whole connectome (Shen *et al.*, 2017). Since the 100-component basis set performed slightly better than the others in cross-validation within the training dataset, we focused on this basis set for comparison with CPM in the held out test set.

For each of the 11 phenotypes, we trained both methods in the training dataset and tested accuracy of phenotypic prediction in the held out test dataset. Results showed that performance of BBS was comparable to or better than CPM on all 11 phenotypes (comparable to CPM on 8 phenotypes and better than CPM on the other 3 phenotypes; Table 1).

#### 3.1.4 Role of parcellation density

We next examined the robustness of the preceding three analyses to parcellations of varying densities. We used Craddock et al.’s parcellations derived from a spectral clustering algorithm with *K*, the prespecified number of parcels, set from 100 to 900 in increments of 100. While there were some differences observed with the most sparse parcellation (*K*=100), for all analyses in which *K* exceeded 200, the results were highly stable and broadly similar to what we observed with the Power parcellation with 264 ROIs (Figure 3).

### 3.2 Network Structure of Components of Cross-Individual Connectome Variation

We next turn to characterizing connectivity patterns in the components themselves. Figure 5, panels A through C, shows the first three components with nodes organized by membership in ICN communities (e.g., default network, fronto-parietal network, etc.) according the node assignments of Power et al. (2011). Qualitatively, these components appear to exhibit prominent ICN structure: the lines on these figures, which represent boundaries of ICN-ICN interrelationships, appear to be highly informative for characterizing connectivity patterns in the components.

**Figure 5:**
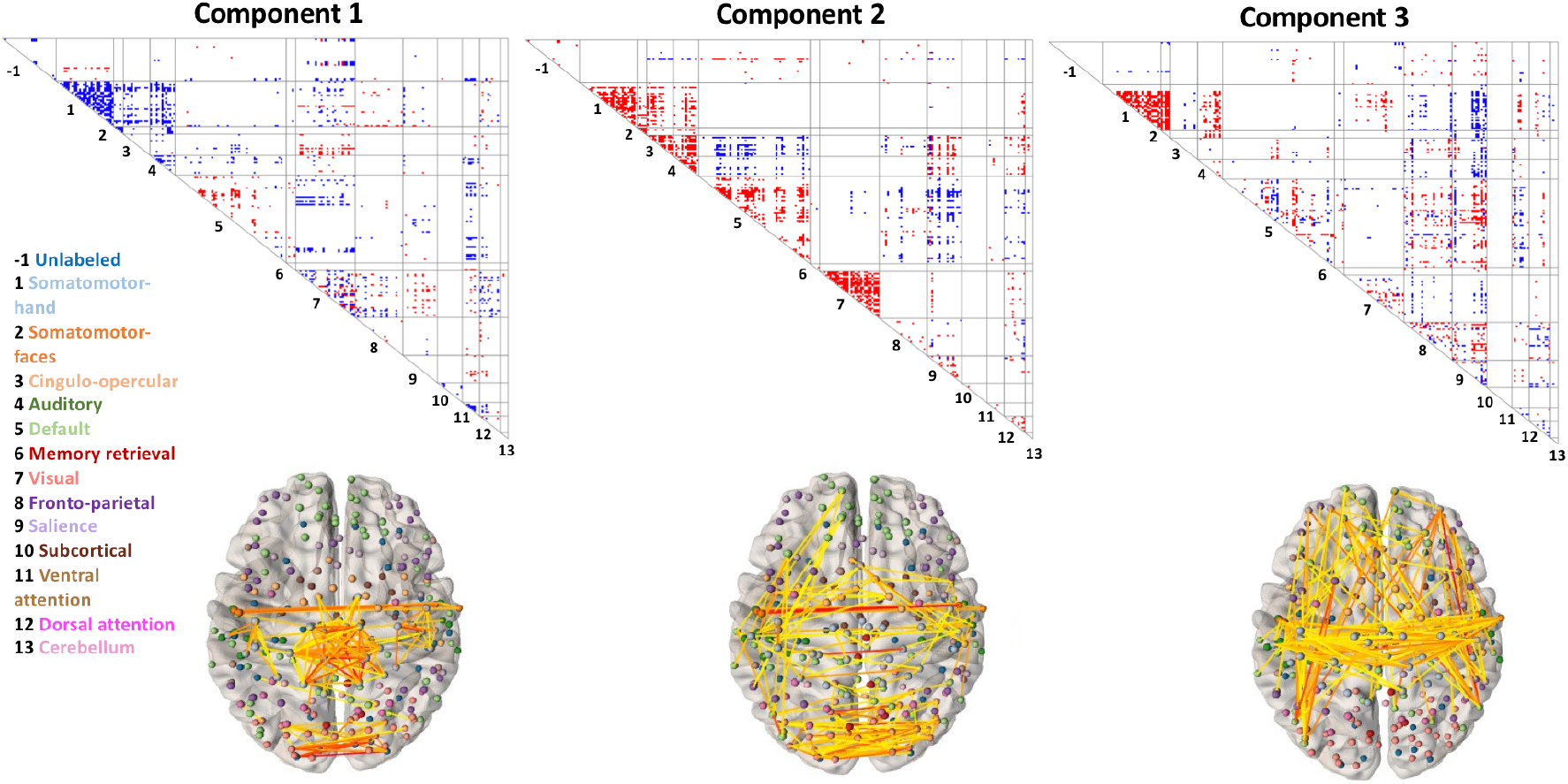
Components 1, 2, and 3. The first three components of inter-individual connectomic variation are displayed, with nodes organized by membership in 13 ICNs according to assignments of Power *et al.*, 2011. The boundaries of ICNs are determined from a strictly intra-individual phenomenon: coherence of the BOLD time series within a person across time. It is notable, then, that inter-individual connectomic differences clearly involve substantial ICN structure (which we further corroborate utilizing a novel quantitative approach based on stochastic block modeling). 1=Somatomotor-hand; 2=Somatomotor-faces; 3=Cingulo-opercular; 4=Auditory; 5=Default; 6=Memory retrieval; 7=Visual; 8=Fronto-parietal; 9=Salience; 10=Subcortical; 11=Ventral Attention; 12=Dorsal Attention; 13=Cerebellum

To quantitatively assess the presence of ICN-based community structure in these components, we utilized an SBM-based method as described in ***Methods*** (see 2.10) coupled with permutation tests for statistical significance. We found that for all 809 components, the observed components’ connectivity patterns are highly statistically significantly more likely under Power ICN community assignments than alternative randomly shuffled assignments (permutation-based p-values for all components survive Bonferroni correction for 809 tests with α = 0.05). Additionally, as a descriptive follow up to quantify the extent of network structure in the components, we investigated how, for each component, the profile log-likelihood corresponding to the Power et al. parcellation differed from the median profile log-likelihood across the permutations (see Figure 6). This analysis suggests that while ICN structure is significantly present in all components, such structure is most prominent in early components and plateaus substantially around component 100 to 200.

**Figure 6:**
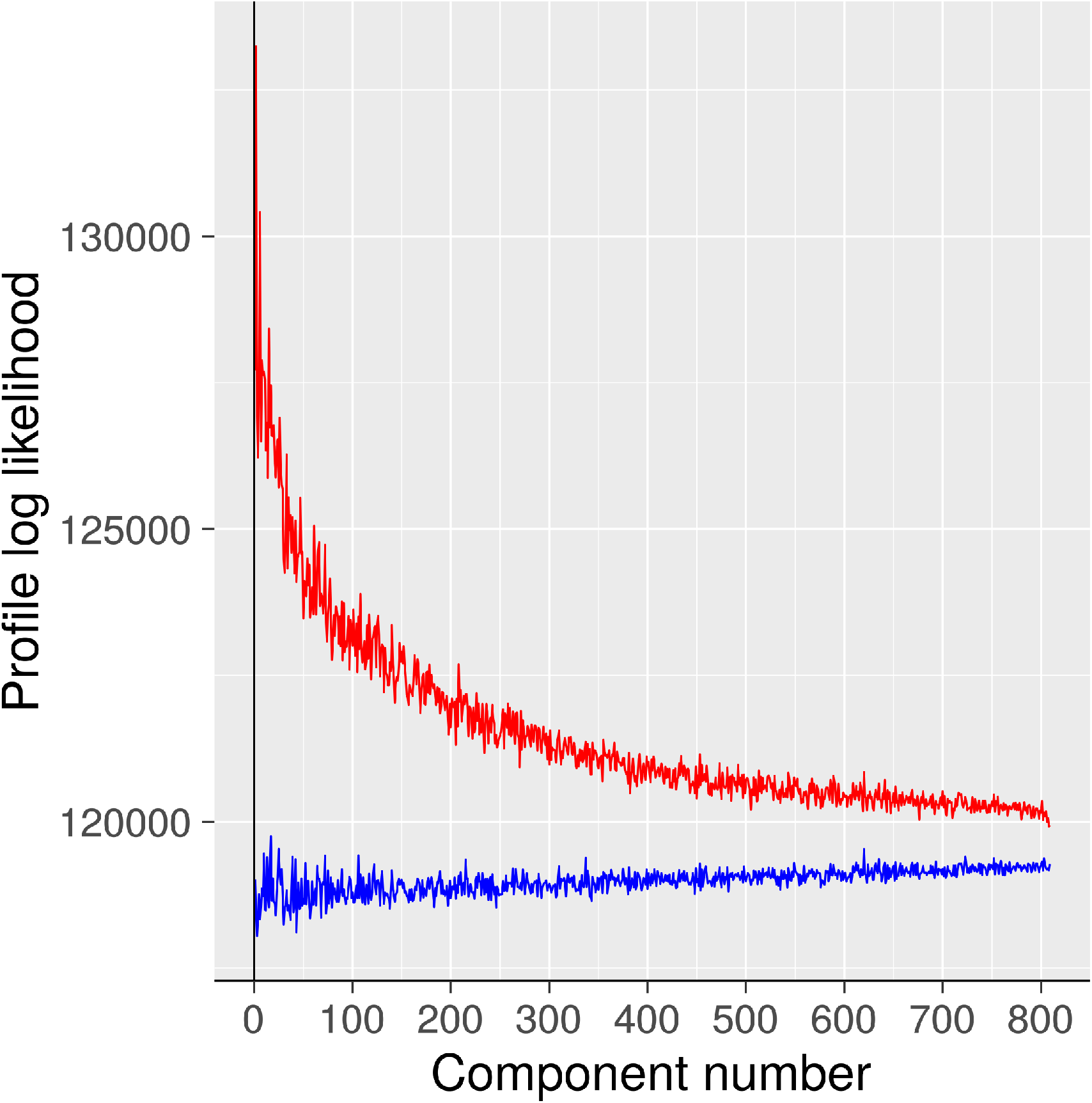
Profile log-likelihood of ICN-based community structure in each component. This statistic serves to quantify presence of ICN structure in each component using a stochastic block model (SBM) framework. The red trace is the profile log-likelihood from the SBM according to the community assignments given in Power *et al.*, 2011. The blue trace is the median profile log-likelihood across many shufflings of the community assignments. See Supplementary Methods for details. ICN structure is most prominent in early (high eigenvalue) component

Given evidence of prominent network structure in the components, especially in earlier components, we sought to further characterize their patterns of network interrelationships. Figure 7 shows network-to-network relationships for the first 150 components. Visual network, DMN, and FPN are especially prominent. Of note, the 150-component basis set is available for viewing and download here: https://sites.lsa.umich.edu/sripada/data/.

**Figure 7:**
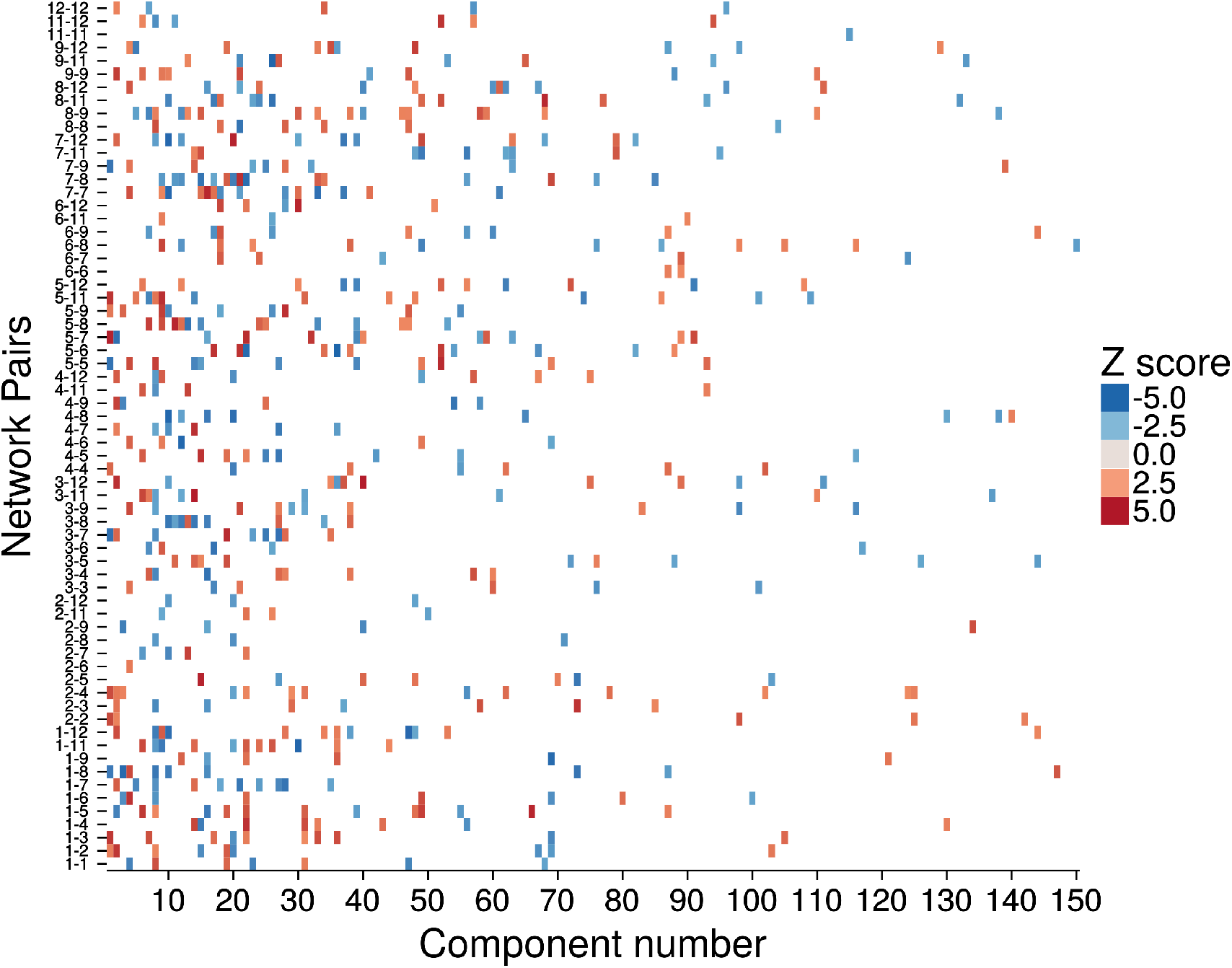
Network Structure of the Connectomic Components. For the first 150 components, altered connectivity patterns are shown using colored squares. Each square represents altered mean connectivity among connections linking pairs of networks. 1=Somatomotor-hand; 2= Somatomotor-faces; 3=Cingulo-opercular; 4=Auditory; 5=Default; 6=Memory retrieval; 7=Visual; 8=Fronto-parietal; 9=Salience; 10=Subcortical; 11=Ventral Attention; 12=Dorsal Attention; 13=Cerebellum.

### 3.3 Test-retest reliability

The preceding analyses suggest that a modest-sized basis set is sufficient to quantify cross-individual variation across the entire connectome, especially the meaningful (i.e., phenotypically predictive) aspects of this variation. A further question concerns the stability of the basis set—or more specifically, subjects’ component expression scores—across scanning sessions.

To address this question, we examined the intra-class correlation (ICC) of component expression scores in the 38 HCP test-retest subjects. Components were generated in the full training dataset, and assessed across the two scanning sessions of the test-retest dataset, which was not used to generate the components. The mean ICC for individual edges is .54, similar to values seen in previous studies (Noble *et al.*, 2017). In contrast, the ICC for the components of cross-individual variation are notably higher. Focusing on the 100-component basis set, which performed well in phenotypic prediction, the mean ICC is 0.78.

Some of this improvement might be due to aggregation itself, as aggregates tend to be more stable than the elements that are aggregated. To test this possibility, we calculated the mean ICC for random permutations of these 100 components (1000 permutations of each component). Mean ICC for permuted components was 0.65, so the boost in ICC seen in the actually observed 100 components is substantially over and above what can be explained by simply aggregating random collections of edges.

## 4 Discussion

In resting state fMRI, the presence of low-rank structure in *intra*-individual variation is well known: a small set of units—ICNs such as DMN and FPN—account for a sizable portion of variation in the BOLD signal across time within a scanning session. In this study, we extend the search for useful low-rank structure to *inter*-individual connectomic variation. We found convergent evidence that a modest number of components, roughly 50-150, capture a sizable share of how the resting state functional connectomes of any two healthy adults differ. Moreover, we found these components exhibit high levels of network community structure, aiding interpretability, and they have very good test-retest reliability. We propose that the connectivity components identified in this study form an effective basis set for quantifying and interpreting systematic inter-individual connectomic differences, and for predicting behavioral and clinical phenotypes.

### The components of inter-individual connectomic variation reflect ICN structure

A remarkable feature of the connectomic components that emerged in this study is that they strongly reflect ICN structure. ICN boundaries are determined from a strictly intra-individual phenomenon: coherence of the resting state blood oxygen level dependent (BOLD) time series across regions within a person during a scanning session (Fox *et al.*, 2005; Power *et al.*, 2011; Yeo *et al.*, 2011). There is no necessity that ICNs should be implicated in across-individual differences in functional connectomes; the set of edges that make individuals different could just have easily have crossed ICN boundaries freely. That is not what we found, however, based both on qualitative observation as well as quantitative assessment.

The finding that there is extensive ICN structure in these components jointly helps to illuminate two issues. First, it helps to explain why we were successful in finding low-rank structure in the first place. Second, it potentially illuminates the mechanisms by which the inter-individual differences we observed arose. Both of these points warrant elaboration.

There is growing understanding of the maturational trajectories of large-scale ICNs and principles by which they take shape. Resting state imaging studies in fetuses suggest at least some important ICNs are in a highly immature state in the fetal brain with weak intra-network connectivity and low levels of network separation (van den Heuvel and Thomason, 2016; Grayson and Fair, 2017; Keunen, Counsell and Benders, 2017). Over the course of childhood to early adolescence, massive changes occur: integration of connections within ICNs (Fair *et al.*, 2008, 2009), segregation of default mode network from attention/control networks (Fair *et al.*, 2007; Anderson *et al.*, 2011; Kessler, Angstadt and Sripada, 2016), and cross-modal linkages in which structural connections co-develop with functional connections (Byrge, Sporns and Smith, 2014; Supekar *et al.*, 2010; Goñi *et al.*, 2014; Betzel *et al.*, 2014). Importantly, there are inter-individual differences in how these developmental changes in ICN-ICN interconnections unfold (Kessler *et al.*, 2014; Kessler, Angstadt and Sripada, 2016; Satterthwaite *et al.*, 2013, 2015).

The overall picture, then, involves highly complex and choreographed developmental processes that shape large populations of interconnections between ICNs. This picture is well suited for explaining why we observed significant low-rank structure in inter-individual variation in connectomes, as such structure necessarily exists if individuals systematically differ at large aggregates of connections. In addition, the model explains why the connectomic components themselves exhibit extensive ICN structure, as the presence of such structure naturally follows if the generative processes that produce inter-individual connectomic differences impart aggregate intra- and inter-ICN alterations.

In short, then, we propose that adult inter-individual connectomic variation— especially the meaningful aspects of this variation that is relevant to explaining neurocognitive and behavioral phenotypes—importantly reflects the legacy of inter-individual differences in ICN development. This hypothesis invites detailed future investigation, ideally in longitudinal datasets that permit precise quantification of ICN maturational trajectories as well as adult connectomic variation.

### Success at Phenotypic Prediction and Test-Retest Reliability

The Brain Basis Set (BBS) modeling approach leverages a modest number of components of inter-individual variation—in this study we focused on a 100-component basis set. Yet we found this method predicts HCP phenotypic variables (such as executive functioning, processing speed, and externalizing) just as well, or in some cases better than, Connectome Predictive Modeling (CPM), an alternative highly successful method that is trained on every edge of the connectome (Shen *et al.*, 2017). The most likely explanation for this result is that systematic connectomic differences across individuals really do have substantial low-rank structure. Thus restricting one’s predictor set to a modest number of connectomic components, which is sufficient to capture this structure, yields strong phenotypic prediction.

An additional complementary explanation emphasizes the issue of signal-to-noise ratio and test-retest reliability. While the inter-session test-retest reliability of individual edges of the resting state functional connectome has been found to be only fair (Birn *et al.*, 2013; Noble *et al.*, 2017), the connectomic components identified in this study exhibit substantially better reliabilities. This improvement arises, most likely, because high eigenvalue components—i.e components that explain a large portion of inter-individual connectomic variation—are more likely to be latching onto “real” brain differences, i.e stable cross-individual differences that genuinely exist in nature. In contrast, connectivity features that explain only a tiny portion of inter-individual variation have a greater probability of reflecting noise, which, by definition, lacks test-retest reliability. It follows that restricting analysis to a modest number of high eigenvalue components can boost the signal-to-noise ratio of the included predictors, contributing to better prediction of unseen data (see Amico and Goñi, 2018 for a related argument).

### Uniqueness of the Basis Set

Our primary result concerns the size of the basis set needed to capture meaningful inter-individual connectomic differences. Resting state connectomes, due to their massive size, allow for correspondingly massive variability: individuals could potentially differ in countless ways across tens of thousands of connections. We have shown, however, that actual inter-individual variability is far more limited and most of it is accounted for by a modest-sized basis set of roughly 50-150 components.

We wish to emphasize that with respect to representing the subspace of variation, the connectomic components we identified are not unique. These components are the basis of a subspace, and any rotation that preserves their linear independence will result in a new basis that spans the exact same subspace. There is thus some flexibility in choosing the components with which to characterize the relevant subspace. Ultimately, the choice of which components to utilize must be guided by consilience with broader theory: a basis set should be preferred to the extent that the components that comprise it align with known neurobiological mechanisms and processes. In this context, it bears notice that the PCA-derived components that emerged in this study do exhibit a number of neurobiologically interesting properties. High eigenvalue components, in particular, disproportionately contribute to phenotypic prediction (Figure 3), and they exhibit higher levels of ICN structure (Figure 6). This provides initial evidence that that the specific components found by PCA could potentially have neurobiological meaning.

### Implications for connectomic statistical analysis

Our results have broader implications for methods of statistical analyses of connectomes, especially methods aimed at predicting phenotypic differences across individuals and between groups (Meskaldji *et al.*, 2013; Varoquaux and Craddock, 2013). A persistent challenge in individual differences research has been the shear size of functional connectomes (Zalesky *et al.*, 2012). This sometimes forces researchers to choose between focusing on a small set of “connections of interest” or else undertake a whole connectome statistical search and pay a substantial price in terms of multiple comparisons correction. Our results suggest that the tradeoffs need not be so stark. There is a massive amount of dependence among edges in connectomes across individuals. Thus a basis set with a modest number of components allows researchers interested in individual differences to undertake whole-connectome inquiry while dramatically reducing the multiple comparison cost.

More broadly, there is a pressing need to leverage prior knowledge about the nature, kind, and extent of inter-individual variation in functional connectomes to further guide and constrain statistical models in neuroimaging individual differences research. Our observation of extensive low-rank structure, i.e a modest number of components account for a sizable portion of cross-individual differences, represents one kind of prior knowledge. Our observation of prominent ICN structure within these components, discussed earlier, is also highly relevant in this context. Future studies should leverage this observation, for example using block structure-based regularization, to inform and constrain statistical models of inter-individual differences, and thereby increase the chances of robust out-of-sample generalization.

In sum, in this study, we identified a parsimonious basis set for inter-individual differences in resting state functional connectomes, one that facilitates interpretation of connectomic differences and prediction of phenotypes of interest. Our results invite further research into the neurodevelopmental processes that shape ICNs, which could help to explain why adult inter-individual connectomic differences take a modest set of characteristic forms.

## Acknowledgments

CS was supported by R01MH107741 and U01DA041106. CS and LL were supported by a grant from the Dana Foundation David Mahoney Neuroimaging Program. LL and YK were supported by NSF grant DMS-1521551. LL and DK were supported by NSF grant DMS-1646108. Thanks to Tal Yarkoni for useful comments on an earlier draft.

## Author Contributions

Conceptualization: CS, MA, DK; Methodology: CS, LL, MA, YK, DK; Formal Analysis: CS, MA, SR, YK, DK, MY; Data Curation: MA, SR; Writing – Original Draft: CS; Writing – Reviewing and Editing; CS, MA, SR, LL, DK; Visualization: MA, SR, DK; Supervision: CS, LL; Funding Acquisition: CS, LL.

## 1 Details Regarding Assessment of Community Structure

### 1.1 The Stochastic Block Model

The SBM [1] is a well-established generative model for networks with communities. Under the SBM, each of the *n* nodes is independently assigned to one of *K* communities, with probability of assignment to community *k* given by *π_k_*, 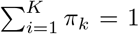. Given a realization of the community assignments vector *c*, where *c_i_* is the community label of node *i*, the SBM generates edge weights *A_ij_* between nodes *i* and *j* independently, from a distribution depending only on the community labels *c_i_* and *c_j_*. If the distribution is parameterized by a parameter *θ_c_i_,c_j__*, the distribution of the entire network is determined by the set of parameters *θ_kl_*, *k, l* = 1 *,…, K*, with *θ_kl_* = *θ_lk_* if the network is symmetric, as it is in our case. In the classical formulation of the SBM, the adjacency matrix is assumed to be binary, in which case the distribution of *A_ij_* is Bernoulli and *θ_kl_* = *P*(*A_ij_* = 1|*c_i_* = *k*, *c_j_* = *l*). In our case, because we work with weighted matrices and the weights are Fisher-transformed correlations, we model the distribution of *A_ij_* as normal, determined by parameters 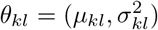.

### 1.2 Calculating profile likelihood under the SBM

In our setting, we have an a priori community membership as given by the Power *et al.* parcellation [2]. The log-likelihood of the observed weights for a given community assignment, *c*, is given by

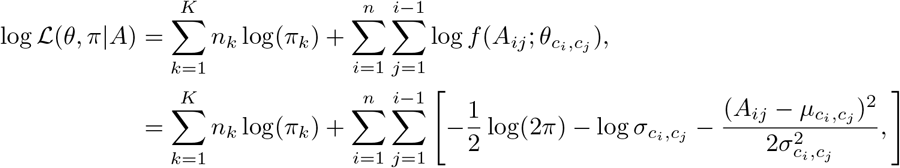

where *n_k_* is the number of nodes in community *k*, and *f*(*·*; θ_kl_) is the probability density function of 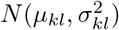.

Maximizing the likelihood of the SBM over community assignments is an NP-hard problem, but for a given *c*, maximizing over *π* and *θ* is easy and there is a closed form solution. Let *S_kl_* denote the set of node pairs connecting community *k* to community *l*, *S_kl_* = {*i < j*: *c_i_* = *k*, *c_j_* = *l*}, and let *n_kl_* = |*S_kl_ |* denote the number of such pairs. Then the maximum likelihood estimates of parameters for a given *c* are

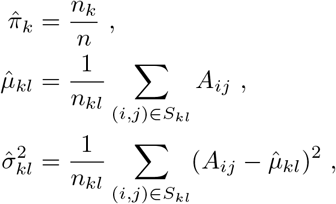

the usual MLEs under the normal distribution. Plugging in these values into the profile likelihood gives the maximized profile likelihood, which we use as the test statistic.

To carry out the test, we need to compare the value of the observed profile log-likelihood, 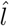, to the distribution of profile log-likelihoods under the null hypothesis of no community structure in the data. We obtain this distribution empirically, shuffing the labels of the given parcellation *c* randomly and recomputing the profile log-likelihood in the same way, *m* = 20,000 times in total, to obtain the values *l_j_*, *j* = 1*,…, m*. Finally, we estimated empirically the probability that a profile log-likelihood *L* sampled from this null distribution will exceed 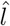, as

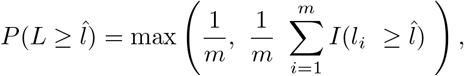

where *I* is the indicator function.

Note that permutation of the labels does not change the number of nodes in each community, so the terms involving 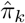’s can omitted.

This procedure is repeated for each of the 809 components of interest, and the resulting 809 p-values are Bonferroni-corrected for multiple comparisons. The number of permutations was selected such that it would be mathematically possible to achieve Bonferroni-corrected significance at α =. 05.

In addition, for each component we retained both the profile log-likelihood under the Power *et al.* parcellation [2] and the median profile log-likelihood across the *m* shuffings, and plotted these as a function of the component number (see Figure XXX). Because of the use of logs, the ratio of likelihoods is proportional to the di↵erence of log-likelihoods, and one may descriptively in-terpret the “gap” between the two traces as some indication of the magnitude of the divergence from the null.

---

**Table S1:** Pearson’s correlations between actual and predicted phenotypes across several predictive models. *BBS = Connectome Basis Set, number following hyphen indicates number of components in basis set; CPM = Connectome Predictive Modeling, pos = positive edges, neg = negative edges (Shen et al. 2017)*.

| ***Phenotype*** | **BBS-50** | **BBS-100** | **BBS-150** | **CPM Pos** | **CPM Neg** | **BBS-100**<br>**covariate**<br>**corrected** |
| --- | --- | --- | --- | --- | --- | --- |
| *General Executive* | 0.43 | 0.44 | 0.37 | 0.42 | 0.32 | 0.39 |
| *Processing Speed* | 0.17 | 0.39 | 0.33 | 0.23 | 0.24 | 0.43 |
| *Penn Progressive*<br>*Matrices* | 0.31 | 0.30 | 0.23 | 0.32 | 0.30 | 0.23 |
| *ASR Externalizing* | 0.20 | 0.24 | 0.17 | 0.03 | 0.06 | 0.25 |
| *ASR Internalizing* | 0.15 | 0.20 | 0.15 | 0.04 | -0.04 | 0.19 |
| *ASR Attention* | 0.14 | 0.21 | 0.07 | 0.00 | -0.02 | 0.20 |
| *NEO-Openness* | 0.14 | 0.18 | 0.23 | 0.11 | 0.07 | 0.14 |
| *NEO-*<br>*Conscientiousness* | 0.17 | 0.19 | 0.16 | 0.15 | 0.13 | 0.11 |
| *NEO-Extroversion* | 0.20 | 0.14 | 0.12 | 0.04 | 0.15 | 0.18 |
| *NEO-Agreeableness* | 0.19 | 0.19 | 0.26 | 0.10 | 0.06 | 0.08 |
| *NEO-Neuroticism* | -0.02 | -0.01 | -0.03 | 0.05 | -0.01 | -0.09 |

